# Nilotinib, an approved leukemia drug, inhibits Smoothened signaling in Hedgehog-dependent medulloblastoma

**DOI:** 10.1101/588897

**Authors:** Kirti K. Chahal, Jie Li, Irina Kufareva, Milind Parle, Donald L. Durden, Robert J. Wechsler-Reya, Clark C. Chen, Ruben Abagyan

## Abstract

Dysregulation of the seven-transmembrane (7TM) receptor Smoothened (SMO) and other components of the Hedgehog (Hh) signaling pathway contributes to the development of cancers including basal cell carcinoma (BCC) and medulloblastoma (MB). However, SMO-specific antagonists produced mixed results in clinical trials, marked by limited efficacy and high rate of acquired resistance in tumors. Here we discovered that Nilotinib, an approved inhibitor of several kinases, possesses an anti-Hh activity, at clinically achievable concentrations, due to direct binding to SMO and inhibition of SMO signaling. Nilotinib was more efficacious than the SMO-specific antagonist Vismodegib in inhibiting growth of two Hh-dependent MB cell lines. It also reduced tumor growth in subcutaneous MB mouse xenograft model. These results indicate that in addition to its known activity against several tyrosine-kinase-mediated proliferative pathways, Nilotinib is a direct inhibitor of the Hh pathway. The newly discovered extension of Nilotinib’s target profile holds promise for the treatment of Hh-dependent cancers.

## Introduction

Smoothened (SMO) is a component of the Hedgehog (Hh) signaling pathway which plays a fundamental role in normal embryonic development and postembryonic tissue homeostasis in eukaryotes^1^. SMO is a seven transmembrane (7TM) receptor that belongs to the Frizzled family^2^. SMO is normally suppressed by a 12TM receptor called Patched (PTCH1) and the suppression is lifted by binding of Patched to a secreted protein ligand Sonic Hedgehog (Shh)^3^. Following release of suppression, SMO translocates from endosomes to the primary cilium where its extracellular cysteine-rich domain (CRD) becomes exposed to a yet unknown natural ligand, proposed to be cholesterol^4^. The activated SMO induces processing of Gli transcription factors into their active form, which subsequently promotes transcription of Gli target genes^1^.

Aberrant Hh pathway activation, both Hh-dependent and independent, is found in several cancers that cumulatively account for about a quarter of all cancer deaths^5^, including basal cell carcinoma (BCC)^6^, medulloblastoma (MB)^7^, myeloid leukemia^8^, rhabdomyosarcoma^9^, pancreatic adenocarcinoma^9^, glioblastoma multiforme (GBM)^10^ and other cancers. MB, a deadly cancer that accounts for 33% of all pediatric brain tumors, is driven by dysregulation of the Hh pathway in about one-third of the cases, and in *majority* of cases in children below the age of five^11^: this MB subtype is referred to as Hh-MB.

Hh pathway is also important in maintenance of cancer stem cells (CSCs), a subpopulation of cancer cells that enable tumor persistence, heterogeneity, and the capacity to self-renew^12^. CSCs are often resistant to chemo- and radio-therapy, which is one of the reasons for tumor resistance and recurrence^13,14^. Because the inhibition of the Hh pathway in CSCs may sensitize these cells to cytotoxic drugs and radiation^12^, the therapeutic relevance of such inhibition may extend beyond those cancers that dysregulate SMO or other components of the pathway in bulk of the tumor.

Among tumors with dysregulated Hh pathway signaling, some are sensitive to SMO antagonists, making SMO a promising anti-cancer therapeutic target^15,16^. Cyclopamine, a naturally occurring teratogenic alkaloid, was identified as the first selective SMO antagonist using cyclopamine derivatives (^125^I-labeled PA-cyclopamine and BODIPY-cyclopamine), and was shown to selectively inhibit Hh pathway activity^17^. Three SMO antagonists were recently approved by the US FDA, Vismodegib (Erivedge^®^) in 2012 for BCC, Sonidegib (Odomzo^®^) in 2015 for BCC and Glasdegib (Daurismo^TM^) in 2018 for acute myeloid leukemia (AML). Several other SMO antagonists are in clinical trials for various types of cancers^16^. Vismodegib, Sonidegib and LY2940680 are currently being actively studied as targeted therapeutics against Hh-MB^18^. Despite the initial promise, the SMO-specific antagonists are often found to be ineffective or to become ineffective over the course of treatment^19^. Therapeutic failure may be caused by escape mutations in SMO^20^ and other components of the Hh pathway^19^, or compensatory changes in other pathways^21^ and cross-talk between different pathways^22^. As a result, only a fraction of Hh-MB patients respond well to the SMO antagonists^23^, and acquired drug resistance or cancer relapse rates are high^20^. Hence, new therapeutic approaches and ideas are urgently needed.

Recently, the cancer research community has increasingly recognized the value of simultaneous targeting of several cancer-related pathways^24,25^. Unfortunately, combination therapies are often poorly tolerated because of disproportional increase in toxicity when several drugs are co-administered. Here we promote an alternative strategy: rather than combining two or more pathway-specific drugs, we propose to look for *useful multi-target profiles of individual drugs* matching a specific cancer subtype. Given the inherent variability of cancers and their escape pathways, this strategy holds the biggest promise when applied in a patient-specific manner^27^. In the context of this strategy, the discovery of realistic multi-target profiles of drugs is particularly important.

To apply this strategy to the Hh-dependent cancers, we searched for anti-SMO activities of existing approved or withdrawn drugs, with a specific focus on drugs with known activity against other cancer-related targets^28^. Using the crystal structures of the transmembrane (TM) domain of SMO^29^, *in silico* structure-based molecular docking^30–32^, and *in vitro* experiments, we identified and confirmed Nilotinib, an approved second generation protein tyrosine kinase inhibitor discovered in 2005^33^, as a potent SMO antagonist. Consistent with this finding, Nilotinib inhibited viability of two Hh dependent MB cell lines (MB-PDX and DAOY) in neurosphere culture, both within clinically relevant concentration range. Nilotinib also reduced tumor volume in a mouse MB xenograft model, and suppressed Gli-1 mRNA in both *in vivo* and *ex vivo* tumor cells. This finding extends the already diverse target profile of Nilotinib (including protein tyrosine kinases BCR-ABL, PGDFR, c-Kit, MK11 and many others)^28,34^ and provides a rationale for using the drug in matching Hh-dependent cancers.

## Results

### *In silico* prediction of compound binding to SMO

As the first step, we set out to identify currently unknown anti-SMO activities of approved drugs using *in-silico* methods and primarily focusing on drugs with established activities against complementary cancer-related pathways. The Drugbank database of approved and withdrawn drugs (together 1699 drugs) was filtered by the logP and Polar Surface Area (PSA) properties to match those of existing SMO antagonists (13 compounds, Supplementary Figure 1) resulting in a dataset of 848 drugs (Figure 1a). Two types of three-dimensional (3D) docking models were employed for drug screening: ligand-based and pocket-based, focusing in both cases on the TM domain of the receptor^29,35^ rather than on its extracellular CRD^4^. Ligand-based 3D atomic property field (APF) models^36^, also referred as chemical field models, were prepared from characterized and co-crystalized ligands of SMO: Cyclopamine, ANTA XV, LY2940680, SAG and SANT-1 (Figure 1b). The pocket docking models for SMO were prepared from multiple Protein Data Bank (PDB) structures of the SMO TM domain (Figure 1c) described in Methods. The 848 drugs along with the 13 known SMO modulators were screened against the ligand-APF models and the pocket docking models to prioritize hits for experimental validation (Figure 1a). Table 1 shows the docking scores and percentile ranks of known SMO modulators and drugs chosen for experimental validation. Vismodegib was the top-scoring compound, SAG and several other known SMO modulators scored well, while Cyclopamine and a closely related drug Saridegib scored poorly. The two poor scores might have resulted from a low resolution (3.2Å) and ambiguities of the available co-crystal structure of Cyclopamine with the TM domain of SMO, as difficulties of docking and scoring Cyclopamine and Saridegib to that site have been reported previously^37^. It has also been suggested that the primary binding site of cyclopamine is located outside of the TM domain, in the CRD of the receptor^4,38^. Overall, eight out of thirteen known SMO compounds were placed in the top 5% of the hit list ordered by the predicted docking scores. These data suggested that the *in silico* screening strategy used in this study is predictive of SMO binding capability of different compounds even though some compounds and modes of binding may be missed.

**Figure 1:**
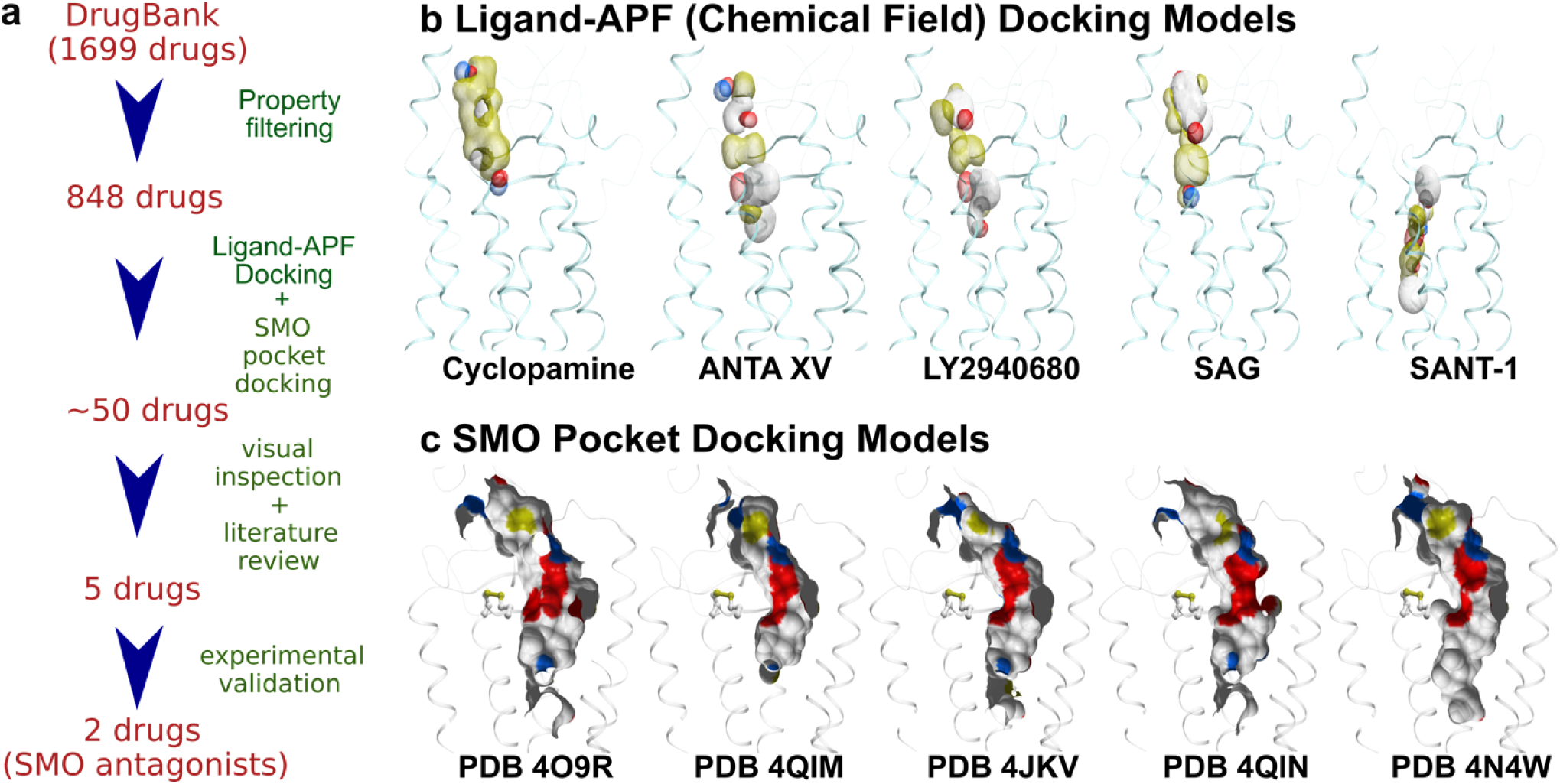
In silico screening pipeline for SMO binders. **(a)** Schematic representation of the procedure **(b)** Ligand-APF Models used for screening of the DrugBank Database. Aromatic and aliphatic features are represented by white and yellow, respectively; hydrogen bond donors and acceptors are blue and red, respectively; positive and negative charges are gray and pink, respectively. **(c)** Pocket Docking Models used for the docking procedure. Carbon, oxygen, nitrogen, and sulfur atoms in the pockets are colored white, red, blue, and yellow, respectively.

**Table 1:**
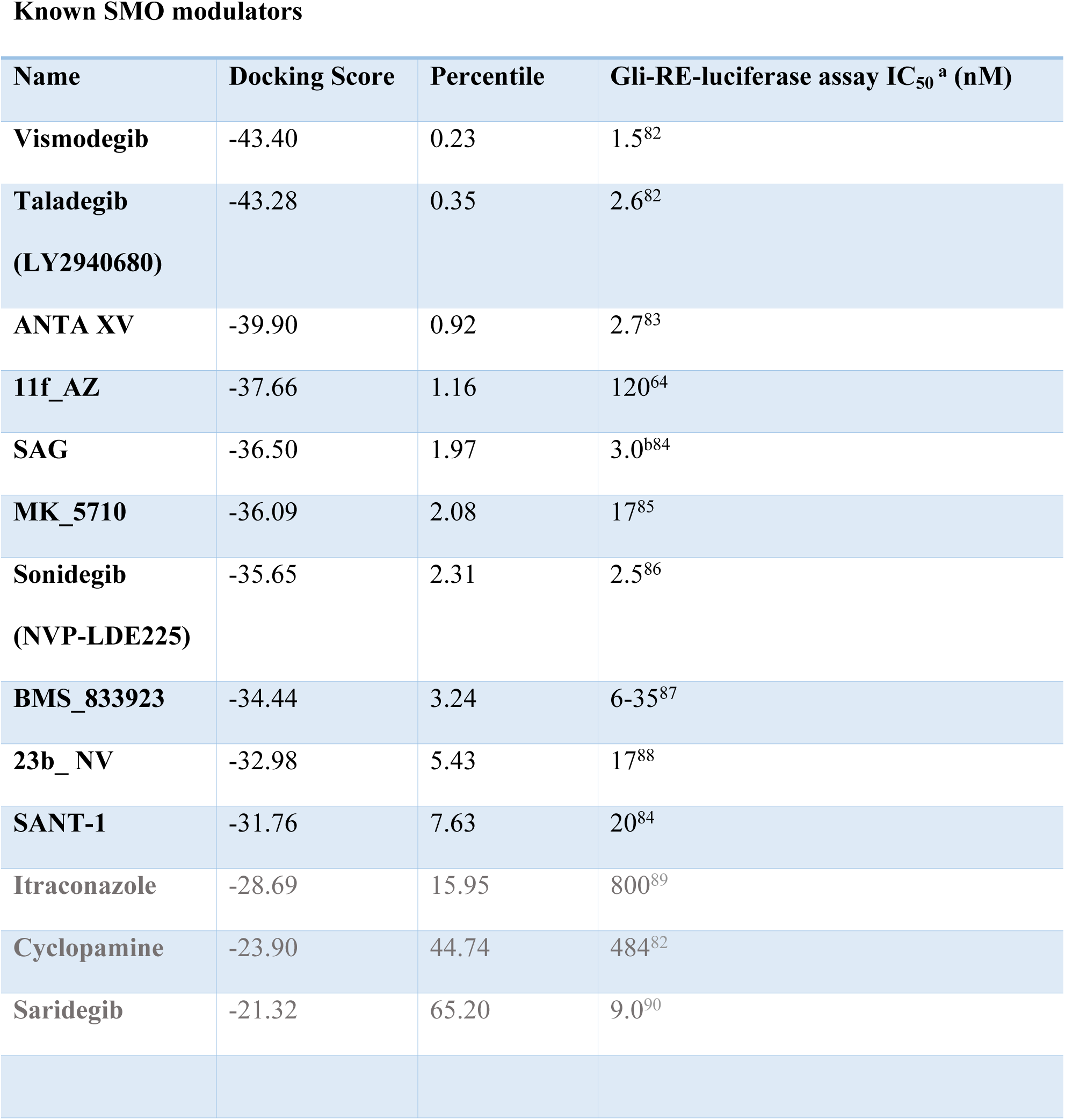
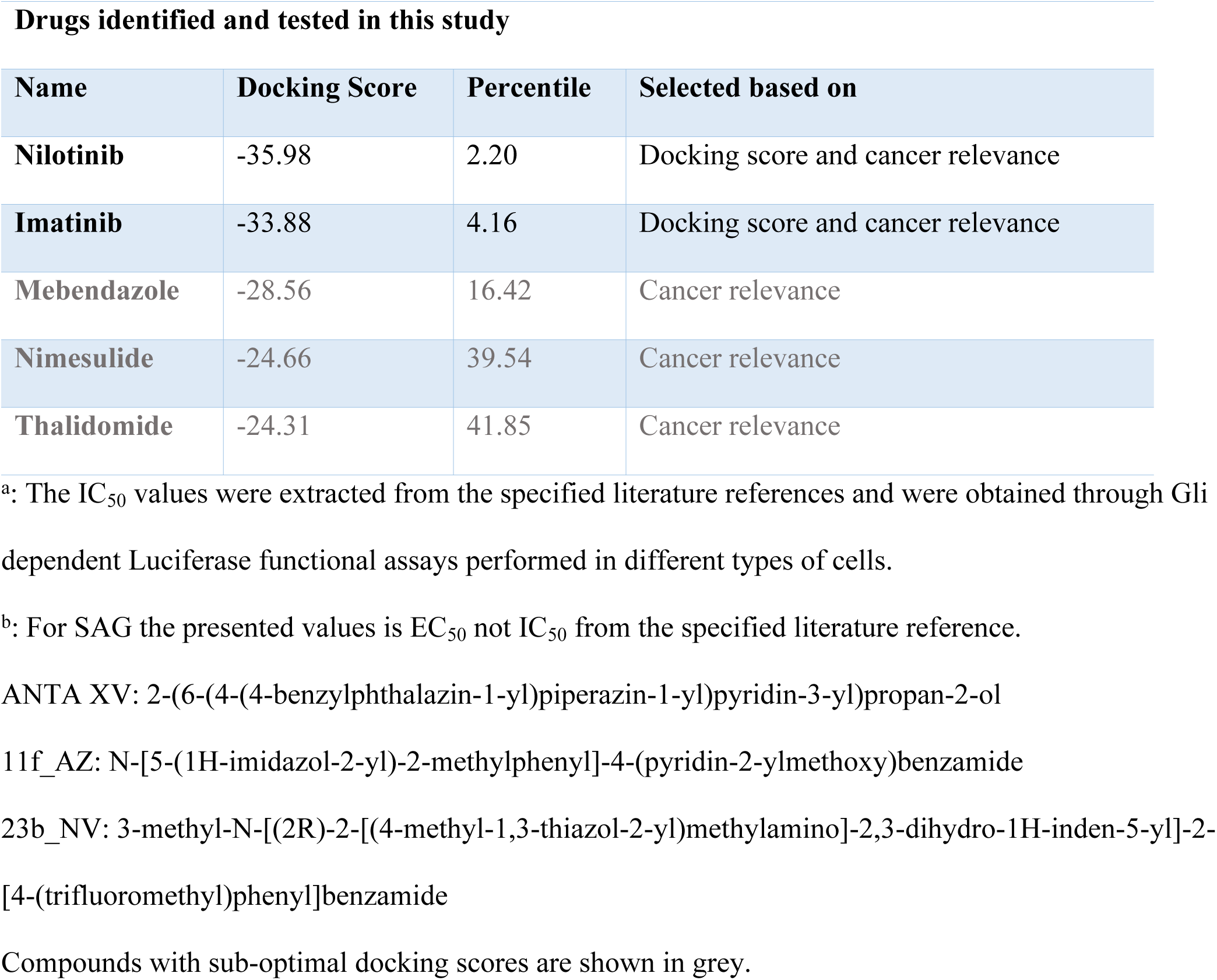
Pocket docking scores, and hit list percentile ranks for known SMO modulators and the selected screening candidates. For the known SMO modulators, literature-reported SMO inhibitory potency is given.

Known SMO modulators
| Name | Docking Score | Percentile | Gli-RE-luciferase assay IC<sub>50</sub><sup>a</sup> (nM) |
| --- | --- | --- | --- |
| **Vismodegib** | -43.40 | 0.23 | 1.5<sup>82</sup> |
| **Taladegib**<br>**(LY2940680)** | -43.28 | 0.35 | 2.6<sup>82</sup> |
| **ANTA XV** | -39.90 | 0.92 | 2.7<sup>83</sup> |
| **11f_AZ** | -37.66 | 1.16 | 120<sup>64</sup> |
| **SAG** | -36.50 | 1.97 | 3.0<sup>b84</sup> |
| **MK_5710** | -36.09 | 2.08 | 17<sup>85</sup> |
| **Sonidegib**<br>**(NVP-LDE225)** | -35.65 | 2.31 | 2.5<sup>86</sup> |
| **BMS_833923** | -34.44 | 3.24 | 6-35<sup>87</sup> |
| **23b_NV** | -32.98 | 5.43 | 17<sup>88</sup> |
| **SANT-1** | -31.76 | 7.63 | 20<sup>84</sup> |
| **Itraconazole** | -28.69 | 15.95 | 800<sup>89</sup> |
| **Cyclopamine** | -23.90 | 44.74 | 484<sup>82</sup> |
| **Saridegib** | -21.32 | 65.20 | 9.0<sup>90</sup> |

Drugs identified and tested in this study
| Name | Docking Score | Percentile | Selected based on |
| --- | --- | --- | --- |
| **Nilotinib** | -35.98 | 2.20 | Docking score and cancer relevance |
| **Imatinib** | -33.88 | 4.16 | Docking score and cancer relevance |
| **Mebendazole** | -28.56 | 16.42 | Cancer relevance |
| **Nimesulide** | -24.66 | 39.54 | Cancer relevance |
| **Thalidomide** | -24.31 | 41.85 | Cancer relevance |

<sup>a</sup>: The IC<sub>50</sub> values were extracted from the specified literature references and were obtained through Gli dependent Luciferase functional assays performed in different types of cells.
<sup>b</sup>: For SAG the presented values is EC<sub>50</sub> not IC<sub>50</sub> from the specified literature reference.
ANTA XV: 2-(6-(4-(4-benzylphthalazin-1-yl)piperazin-1-yl)pyridin-3-yl)propan-2-ol
11f\_AZ: N-[5-(1H-imidazol-2-yl)-2-methylphenyl]-4-(pyridin-2-ylmethoxy)benzamide
23b\_NV: 3-methyl-N-[(2R)-2-[(4-methyl-1,3-thiazol-2-yl)methylamino]-2,3-dihydro-1H-inden-5-yl]-2-[4-(trifluoromethyl)phenyl]benzamide
Compounds with sub-optimal docking scores are shown in grey.

### Nilotinib and Imatinib are predicted *in silico* to bind SMO

The ligand-based APF docking and pocket docking of the Drugbank database produced an ordered hit list. Following visual inspection of the binding poses of 50 top-scoring candidates (Supplementary Table 1) and considering the anti-cancer potential of the candidates, five drugs were selected for experimental validation. The top scoring candidates included Nilotinib and Imatinib, two protein kinase inhibitors used to treat chronic myelogenous leukemia^34^; their docking scores were comparable to those of the known SMO modulators Vismodegib and SAG. In addition, the following drugs were selected for testing: Mebendazole, a broad-spectrum antihelmintic also studied as a repurposing candidate for cancer^39,40^, Thalidomide, an immunomodulatory drug used for the treatment of leprosy as well as multiple myeloma and other cancers^41^, and Nimesulide, a COX-2 selective nonsteroidal anti-inflammatory drug, known to inhibit growth of various cancer cell lines^42^. The pocket docking scores of these drugs are given in Table 1, and the predicted binding poses of Nilotinib, Imatinib and Mebendazole are presented in Figure 2, along with the crystallographic poses of Cyclopamine and SANT-1, and the prospectively predicted pose of Vismodegib (this work was done prior to publication of the crystallographic structure of Vismodegib-bound SMO^43^).

**Figure 2:**
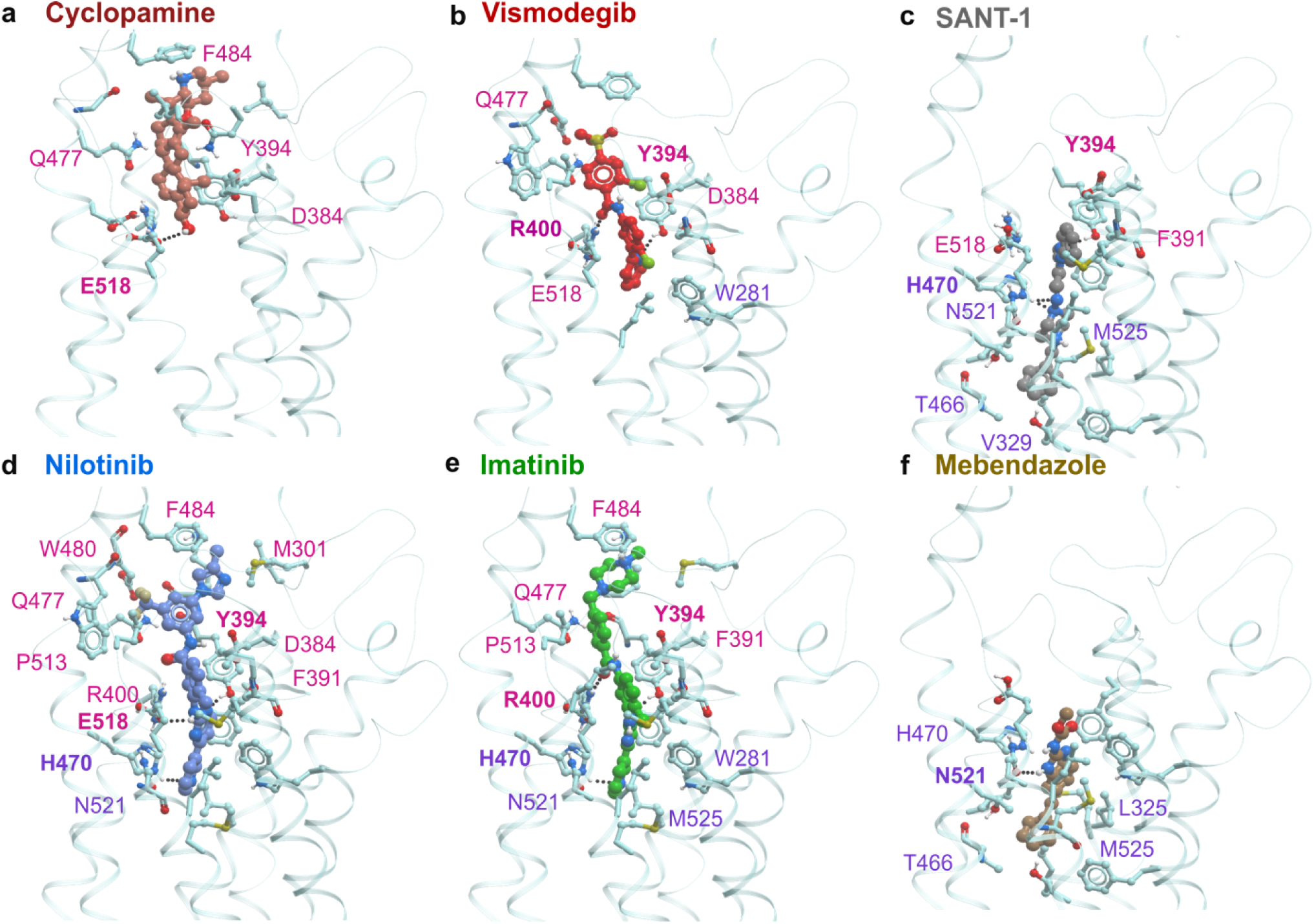
Crystallographic or predicted binding poses of selected SMO ligands (Cyclopamine, Vismodegib, SANT-1, Nilotinib, Imatinib and Mebendazole). The binding cavity of SMO can be tentatively separated into two subpockets – upper binding pocket and lower binding pocket. Cyclopamine **(a)**, Vismodegib **(b)**, and SAG (not shown) bind in the upper binding pocket, while SANT-1 **(c)** binds in the lower binding pocket. In their predicted binding poses, Nilotinib **(d)** and Imatinib **(e)** span both upper and lower binding sub-pockets and form hydrogen bonds with Y394, R400 / E518 and H470. By contrast, Mebendazole **(f)** is more similar to SANT-1 because it occupies the lower sub-pocket only. Receptor residues making contacts with the ligands are shown as sticks. Selected residues making the strongest contacts^81^ are labeled and colored pink (upper binding sub-pocket) and purple (lower binding sub-pocket). Residues making hydrogen bonds with the ligand are labeled in bold print.

Finding two related type-II protein kinase inhibitors, Nilotinib and Imatinib, at the top of the hit list was unexpected. Analysis of their predicted docking poses indicates that these drugs bind SMO in a similar manner (Figure 2d, 2e). In contrast to Cyclopamine or SANT-1 that bind exclusively in the top or the bottom part of the elongated SMO binding cavity, respectively, Imatinib and Nilotinib span both sub-pockets. The pattern of hydrogen bonds formed by Nilotinib (Y394, E518 and H470) and Imatinib (Y394, R400 and H470) is similar to the pattern observed in co-crystallized ligands, or the pattern for Vismodegib (Y394 and R400) in its predicted binding pose (Figure 2b). Nilotinib has a more extensive list of residues participating in the binding as compared to Vismodegib, which may be beneficial.

We next analyzed whether the binding cavity of SMO bears any resemblance with the binding site of the primary target of Nilotinib and Imatinib, the protein kinase ABL1^34^. The analysis of two binding cavities (ANTA XV bound to SMO - PDB 4QIM and Nilotinib bound to ABL1 – PDB 3CS9) indicated that the overall shape and composition of both pockets were similar (elongated narrow channels with a large number of polar features, Figure 3), but the geometric details and specific arrangement of functional groups within the two pockets were different. Furthermore, the predicted conformations of Nilotinib and Imatinib in the SMO binding pocket during docking studies (Figure 2d and 2e) were different from those crystallographically observed in the ABL1 binding pocket (Figure 3). Therefore, this predicted anti-SMO activity of the protein kinase inhibitors could not be explained by a trivial pocket similarity and instead resulted from the ability of flexible drugs to adopt diverse conformations matching dissimilar target pockets.

**Figure 3:**
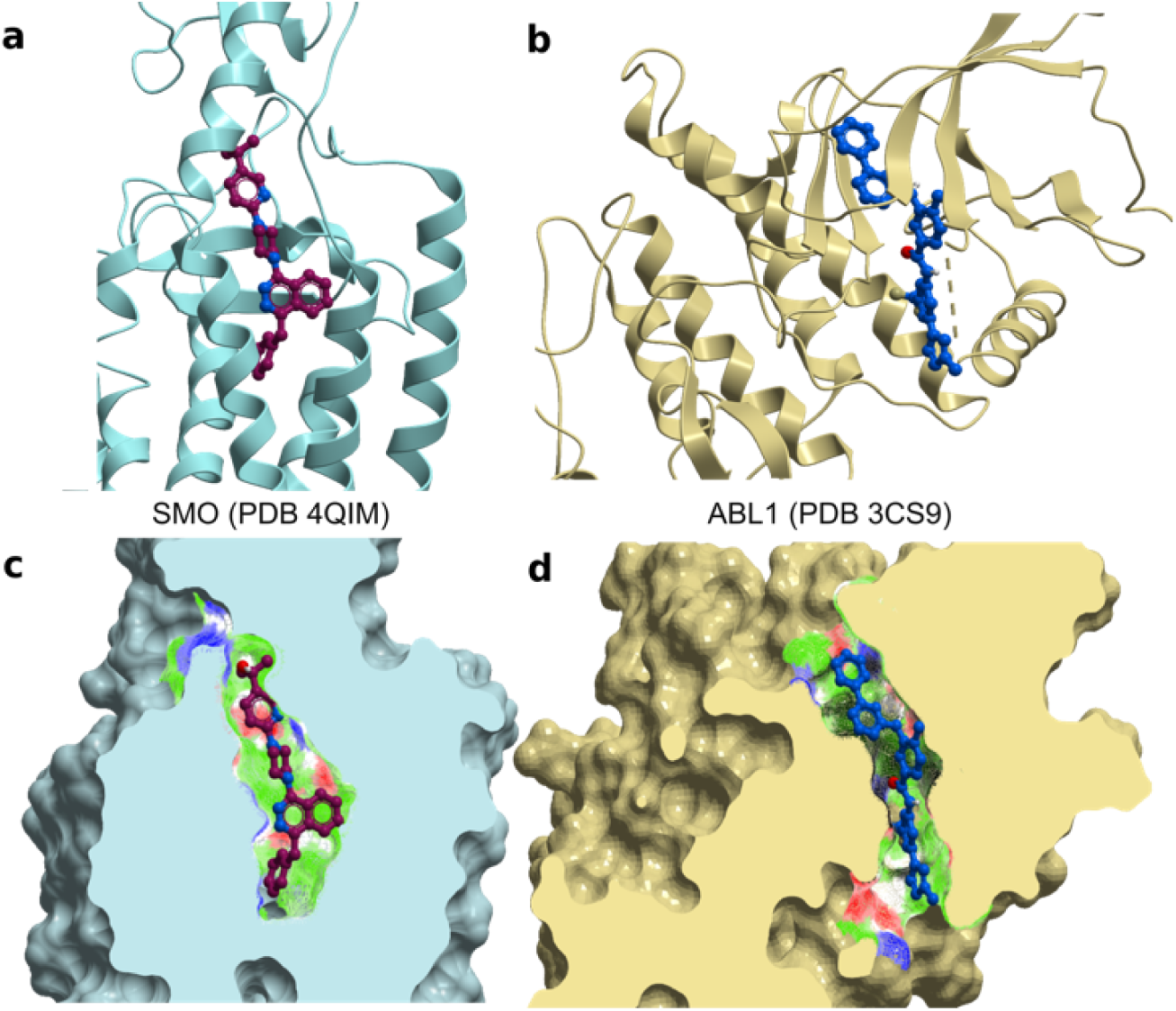
Similarity between SMO and ABL1 binding cavities. **(a)** Ribbon diagram of structure of the SMO bound to ANTA XV antagonist (PDB 4QIM) **(b)** Ribbon diagram of structure of ABL1 bound to Nilotinib (PDB 3CS9) **(c)** Molecular surface of the TM binding pocket of SMO with ANTA XV **(d)** Molecular surface of the binding pocket of ABL1 with Nilotinib.

In contrast to Nilotinib and Imatinib (Figure 2d, 2e), Thalidomide and Nimesulide were predicted to bind to the middle part of SMO binding pocket (not shown). Mebendazole was predicted to bind to the lower sub-pocket of SMO (figure 2f) and to form contacts with the same residues as SANT-1 (Figure 2c and 2f). Interestingly, Mebendazole was independently reported to inhibit the Hh pathway and to reduce growth of Hh-dependent MB cells in an orthotopic xenograft tumor model^44^. Finally, the recently identified experimental dual MET and SMO modulators^28^ were out of scope of this study since we only included FDA approved/withdrawn drugs.

### Nilotinib and Imatinib inhibit Hh-pathway activity

Nilotinib, Imatinib, Mebendazole, Thalidomide, and Nimesulide were tested in a functional Hh pathway activity assay using NIH 3T3 Gli-RE cells stably expressing firefly luciferase under the control of 8x Gli response element (8xGli-RE) and stimulated with the active soluble form of Shh (further referred to as ShhN). Cyclopamine, a previously characterized SMO antagonist^17^, was used as a positive control. Vismodegib, an FDA approved SMO antagonist, was also tested along with other drugs and IC_50_ of 4 nM was observed. In this assay, Nilotinib, Imatinib and Mebendazole inhibited ShhN-induced Hh signaling in a dose-dependent manner. The observed IC_50_ of Cyclopamine was 189 nM, while Nilotinib had IC_50_ of 374 nM (Figure 4 and Table 2). Mebendazole and Imatinib exhibited IC_50_ values of 327 nM and 4.57 μM, respectively (Table 2). Thalidomide and Nimesulide were found inactive, which is consistent with their poor docking scores (Table 1). Neither control nor test compounds produced any effect in the absence of ShhN treatment, indicating that the observed inhibition was specific to the Hh pathway (Supplementary Figure 2). Compounds also had minimal to no effect on luciferase activity independent of SMO expression (Supplementary Figure 3).

**Figure 4:**
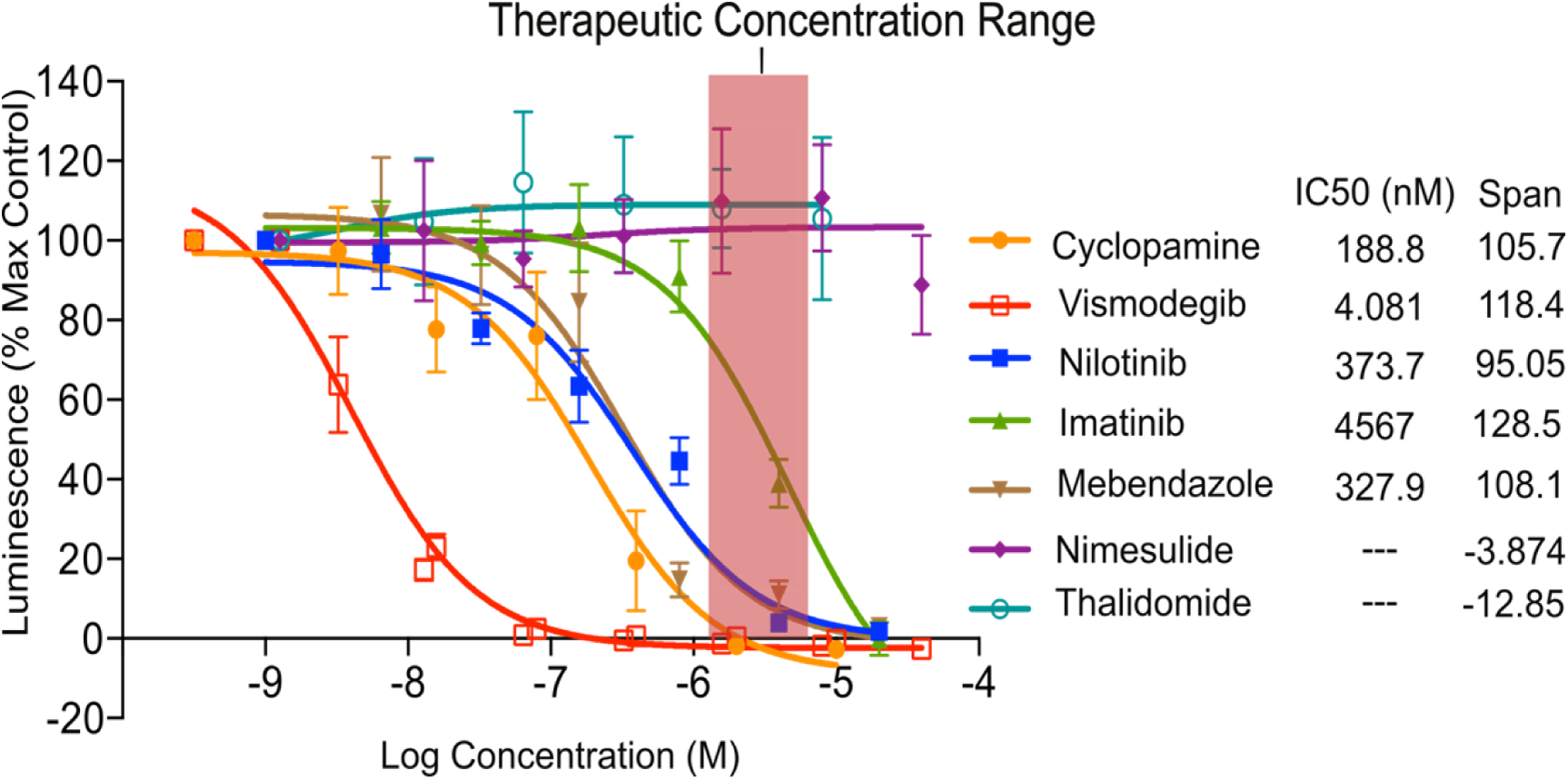
Hedgehog pathway inhibition by Nilotinib, Imatinib and Mebendazole. The drugs were applied at different concentrations to the NIH 3T3 Gli-RE cells followed by stimulation of the cells with ShhN-conditioned media (n=3). The data is presented as percent of the maximum luminescence in the absence of antagonists. Cyclopamine and Vismodegib were included as control antagonists. Nilotinib, Imatinib and Mebendazole, but not Nimesulide or Thalidomide, demonstrated dose-dependent inhibition of ShhN-induced Hh pathway activation. Reported clinically achievable therapeutic concentrations of Nilotinib are marked with a pink rectangle. Data represent the mean and standard deviation.

**Table 2:** IC_50_ values of control and test compounds observed in various assays.

| Name | Hh/SMO pathway<br>inhibition IC<sub>50</sub> (nM)<sup>a</sup> | SMO binding IC<sub>50</sub><br>(nM)<sup>b</sup> | Cell viability assay IC<sub>50</sub> (μM)<sup>c</sup> |  |
| --- | --- | --- | --- | --- |
|  | NIH3T3 cells | HEK293T cells | MB-PDX cells | DAOY cells |
| Cyclopamine | 189 | 123 | ^ | ^ |
| Vismodegib | 4 | 10 | 52 | ~ |
| Nilotinib | 374 | 3105 | 5 | 6 |
| Imatinib | 4567 | 5149 | 17 | 65 |

<sup>a</sup>: IC<sub>50</sub> values were calculated from Gli-luciferase functional assay using NIH3T3 Gli-RE cells. (n=3)
<sup>b</sup>: IC<sub>50</sub> values were estimated from BODIPY-Cyclopamine competition binding assay using HEK293T cells transiently transfected with mSMO. (n=2)
<sup>c</sup>: IC<sub>50</sub> values were calculated from the alamarBlue™ Cell Viability assay. (n=3)
^: Not evaluated
~: Data range not sufficient to derive IC<sub>50</sub> value

The observed functional IC_50_ of Nilotinib favorably compares to clinically achievable plasma concentration of the drug. The trough concentration (C_min_) observed in human plasma in the course of standard dosing of 400 mg twice daily ranges from 1.65 to 2.38 μM^45^ (4.4 to 6.4 times our measured anti-Hh IC_50_). Another study quotes average Nilotinib C_min_ and C_max_ at 2.6 and 3.3 μM^46^, respectively, corresponding to 7 and 8.8 times anti-Hh IC_50_ of 374 nM. These concentrations translate into 81% to 90% inhibition of SMO during Nilotinib treatment. With anti-Hh IC_50_ values of 188 nM and 327 nM (i.e. close to IC_50_ of Nilotinib), Cyclopamine and Mebendazole were both shown to be efficacious in reducing brain tumor size in mice^44,47^. This suggests that Nilotinib may have a similar effect *in vivo*.

### Nilotinib directly binds to SMO TM domain

As demonstrated by both crystallography and biochemistry, SAG and Cyclopamine can directly bind to the 7TM domain of SMO^43,48^. By doing so, they are believed to influence the activation-associated conformational changes in the receptor thus activating or inhibiting the downstream Gli signaling. To check whether the tested compounds bind to SMO at the SAG/Cyclopamine 7TM binding site, competition binding and functional assays were performed. The competition binding was evaluated by flow cytometry in HEK293T cells transiently transfected with SMO, using BODIPY-Cyclopamine as a fluorescent probe. Nilotinib and Imatinib were found to displace BODIPY-Cyclopamine (supplemented at 5 nM) from SMO in a dose-dependent manner, whereas Mebendazole, Thalidomide and Nimesulide showed no effect on BODIPY-Cyclopamine binding (Figure 5b and Table 2). Figure 5a shows the distribution of cell fluorescent intensities of five representative samples stained or not with BODIPY- Cyclopamine with or without competitors. With the exception of Mebendazole, these results are in agreement with the results of the functional assay.

**Figure 5:**
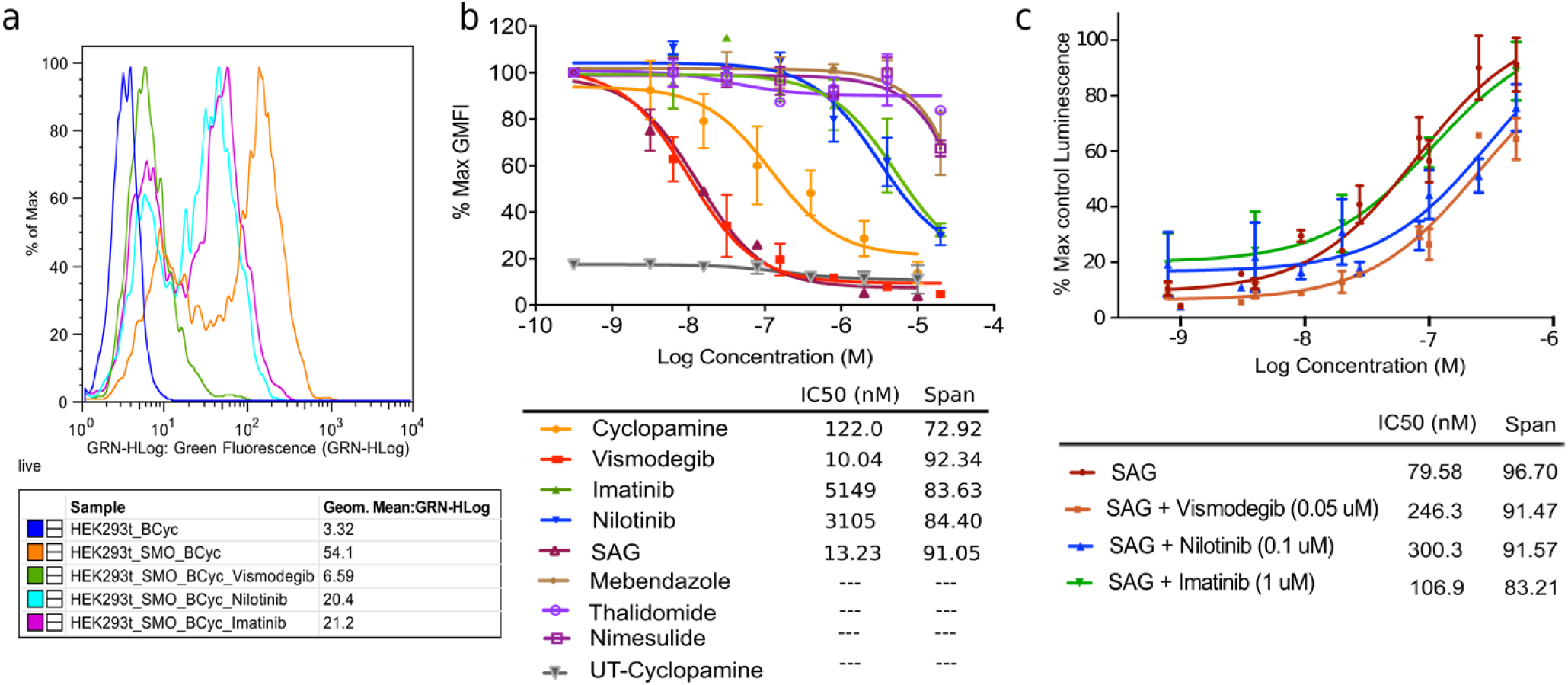
Nilotinib binds to SMO and competes with Cyclopamine and SAG. **(a)** Representative distributions of cell fluorescence intensities measured by flow cytometry and demonstrating competitive displacement of BODIPY-Cyclopamine by Vismodegib, Nilotinib and Imatinib. (**b)** Concentration response curve for flow cytometry based competitive binding assay in HEK293T cells transiently transfected with mSMO: positive controls (Cyclopamine and Vismodegib) and test compounds - Nilotinib and Imatinib displaced BODIPY-Cyclopamine from SMO in a dose-dependent manner. Mebendazole did not demonstrate any appreciable displacement of BODIPY-Cyclopamine in this assay. Data represent the mean and standard deviation for two independent experiments, max GMFI: Maximum Geometric Mean Fluorescence Intensity; UT: Untransfected HEK293T cells (**c)** Concentration response curve of SAG in the Gli-Luciferase functional assay in the absence or presence of inhibitors: Nilotinib or Vismodegib shifted EC_50_ of SAG to the right without decrease in maximal response, suggesting a competitive relationship. Imatinib did not shift EC_50_ of SAG possibly due to its low potency as observed in the functional assay. Data represent the mean and standard deviation for three independent experiments.

In the case of Mebendazole, the discrepancy between the binding and the functional assay results can be explained by the fact that Mebendazole was predicted to bind exclusively in the lower sub-pocket of SMO, whereas Cyclopamine occupies either the upper sub-pocket of the TM domain or even the CRD^4,48^. Therefore, it is conceivable that the two compounds can bind non-competitively and may even form a ternary complex (Figure 2f). In the case of Nilotinib and Imatinib, the assay confirmed their direct binding to SMO; however, the observed binding assay IC_50_ was significantly higher than the functional assay IC_50_ (Table 2). Discrepancies between functional and binding IC_50_ have been observed previously for SMO antagonists and attributed to overlapping but distinct binding sites for the compounds and probes^49^. In particular, for cyclopamine, two distinct binding sites have been demonstrated both biochemically^4^ and crystallographically^4,38,43,48^.

To further explore the mechanism of inhibition of SMO by Nilotinib and Imatinib, we tested it in a competitive functional inhibition assay against SAG, a known small molecule agonist of SMO that binds in the same part of the 7TM pocket^29^ as predicted for Nilotinib. In NIH 3T3 Gli-RE cells, SAG caused robust dose-dependent increase in luminescence with the EC_50_ of 79 nM. When the assay was repeated in the presence of increasing concentrations of Nilotinib, the maximum signaling level induced by SAG was not affected, but its EC_50_ was shifted, indicating competitive inhibition (Figure 5c). Vismodegib, which binds in the same sub-pocket as SAG, was used as a positive control and exhibited similar competitive behavior. Imatinib failed to shift EC_50_ of SAG significantly at the tested concentration, which can be explained by its weak potency. These results further support the predicted binding mode of Nilotinib to SMO in its TM domain pocket.

### Nilotinib reduces nuclear Gli-1 and viability of MB cell lines

To study downstream effects of the Hh pathway inhibition by Nilotinib and other identified drugs, we evaluated their effects on viability and neurosphere (NS) formation capacity of two Hh-dependent MB cell lines: one a patient-derived MB-PDX line^51^ and another an established DAOY line^52,53^. Both MB cell lines are heterogeneous and contain a small sub-population of neural (cerebellar) cancer stem cells among other, differentiated cells. The neural stem cells are capable of forming neurospheres from single cells when seeded sparsely and cultured in non-adherent conditions in serum-free medium supplemented with growth factors. The NS formation assay is commonly used to study the effect of potential therapeutic agents on neural cancer stem cells^54,55^.

To distinguish the effect of drugs on neural cancer stem cells from non-specific cell toxicity, the cell viability measurements were performed side-by-side in the NS favoring conditions (in which only CSCs survive and grow), and in adherent cell culture conditions (in which all cells can grow). Vismodegib was used as a positive control and Erlotinib, an FDA-approved multi-kinase inhibitor^56^ was used as a negative control. When the cells were cultured in adherent conditions in the presence of serum, treatment with drugs did not significantly affect cell viability. By contrast, exposure of cells cultured under NS conditions to Nilotinib significantly reduced both cell viability and neurosphere formation capacity in a dose-dependent manner (Figure 6a - d). Nilotinib was found to be more potent than Imatinib, in agreement with their respective potencies measured in the functional reporter assay. As expected, Erlotinib did not affect cell viability under any condition. Vismodegib did have an effect on cell viability in neurosphere culture conditions, but only at a much higher concentration. The IC_50_ of Nilotinib, Imatinib and Vismodegib were determined to be 5 μM, 17 μM and 52 μM, respectively (Figure 6b, Table 2) in MB-PDX cells. The viability DAOY cells in NS conditions was also affected by Nilotinib: the IC_50_ of Nilotinib and Imatinib were determined to be 6 μM and 65 μM, respectively (Figure 6c, Table 2). The Vismodegib data range was not sufficient to derive the IC_50_ value. The results show that unlike Vismodegib or Erlotinib, Nilotinib can inhibit growth of MB cells significantly at clinically achievable concentrations (trough concentration, C_min_, ranges from 1.6 to 2.6 μM^45,46^).

**Figure 6:**
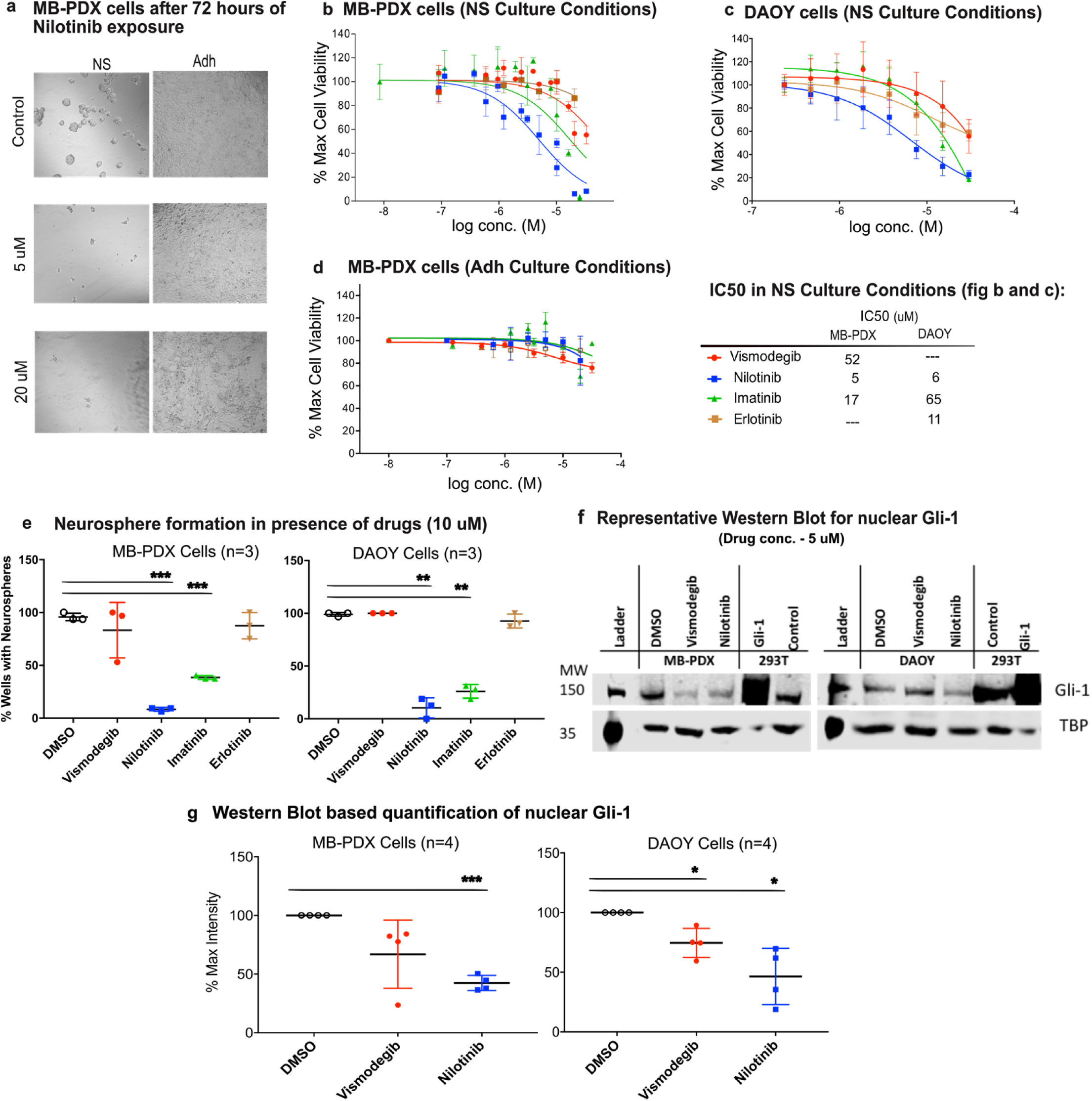
Nilotinib inhibit cell viability and neurosphere formation in medulloblastoma cells (MB- PDX and DAOY). **(a)** Representative bright field images of MB-PDX cells cultured in Neurosphere (NS) and Adherent (Adh) conditions with vehicle and Nilotinib (5 and 20 mM) (magnification 10×). Concentration response curve for cell viability in low serum (NS-formation) conditions (n=3) in MB- PDX cells (**b)** and (**c)** Concentration response curve for cell viability in low serum (NS-formation) conditions (MB-PDX cells – b and DAOY cells – c). The IC_50_ values for cell viability assay observed in MB-PDX and DAOY cells are also provided next to panel (c) under the heading “IC50 in NS Culture Conditions”. (**d)** Concentration response curves for inhibition of cell viability in adherent conditions. Drugs did not affect viability of cells grown in adherent conditions. (**e)** Neurosphere formation assay performed three times in presence of 10 mM concentration of drugs in MB-PDX and DAOY cells. Nilotinib was observed to be most potent in inhibiting the formation of NS from single cells. (**f)** A representative Western blot for quantitation of nuclear Gli-1 in MB-PDX and DAOY cells shows a decrease in Gli-1 protein after treatment with 5 mM Nilotinib for 24 hours. Tata Binding Protein (TBP) was used as loading control for nuclear fractions. For positive controls (mSMO-transfected HEK293T cell lysate), a reduced amount of cell lysate was loaded to avoid overstaining of the blot: this explains disproportional loading controls one sample in each blot. (**g)** Quantification of nuclear Gli-1 in MB- PDX and DAOY cells treated or not with drugs. Image represents the mean and standard deviation of at least three experiments performed on different days. Asterisks represent p-values: *, p < 0.05; **, p < 0.005; ***, p < 0.0005.

In addition to studying cell viability, we also quantified formation of NSs in the sparsely seeded MB- PDX and DAOY cell cultures. Cells were plated at 1–2 cells per well in non-adherent plate and cultured in serum-free medium with growth factors for at least 7 days, in the absence or presence of drugs. Vismodegib and Erlotinib were used as positive and negative controls, respectively. At the concentration of 10 µM, Nilotinib inhibited both MB-PDX and DAOY NS formation in more than 90% of wells (Figure 6e). Imatinib was less potent, and Vismodegib and Erlotinib appeared ineffective (Figure 6e) in this assay.

The fact that Vismodegib appeared much less potent and efficacious than Nilotinib in both cell viability assays and NS formation assays, may be attributed to the differences in mechanisms of action of the two drugs. Vismodegib is a specific inhibitor of SMO, whereas Nilotinib is also capable of inhibiting multiple kinases and related pathways that may be important for cancer cell growth and maintenance^34^. The favorable target profile of Nilotinib may have contributed to the observed effect in these assays. Imatinib shares some features of the multi-target profile of Nilotinib but it is a less potent SMO antagonist. On the other hand, as illustrated by Erlotinib, a multi-kinase profile alone is not sufficient to achieve the desired biological effect: it is a combination of anti-SMO and anti-kinase activity that appears to enable it.

Next, we examined the effects of Vismodegib and Nilotinib on the nuclear expression of Gli-1, the canonical transcription factor activated downstream of SMO, in MB-PDX and DAOY cells cultured in NS conditions. Nuclear Gli-1 was quantified by Western blotting, and HEK293T cells transiently transfected with Gli-1 were used as a positive control. In MB-PDX cells, 24-hour treatment with Vismodegib and Nilotinib reduced mean values of nuclear Gli-1 to 67% and 41% of untreated samples, respectively. Similarly, Vismodegib and Nilotinib reduced mean values of nuclear Gli-1 in DAOY cells to 75% and 46% of untreated samples. Figure 6g shows a representative blot for each of the two cells lines and a cumulative quantitative summary of three independent experiments. The uncropped blots are provided as Supplementary Figure 4. This Western blot analysis confirmed the direct effect of Nilotinib on the Hedgehog pathway through the reduction of nuclear Gli-1, which is similar to or exceeding the reduction caused by Vismodegib. These results are also consistent with the observed superior inhibitory effect of Nilotinib in the cell viability and NS formation assays.

### Nilotinib reduces Gli-1 mRNA and MB-PDX tumor growth *in vivo*

To assess the effects of Nilotinib on tumor growth *in vivo*, human MB-PDX cells^51^ were injected subcutaneously in NSG mice. Two independent experiments were conducted, encompassing the total of 12 tumors in three mice in the control group and 24 tumors in six mice in the Nilotinib-treated group. Nilotinib was administered at 40 mg/kg daily starting from day seven post-grafting. Tumor volumes were monitored three times a week in the course of treatment. After 42 days, all mice were sacrificed, tumors were extracted, measured, and subjected to qRT-PCR.

Nilotinib-treated group developed significantly smaller tumors than the vehicle-treated group (Figure 7a). Moreover, the qRT-PCR analysis of tumors showed a significant reduction in the Gli-1 mRNA expression levels in Nilotinib-treated group in comparison to vehicle treated group (Figure 7b). These data suggest that Nilotinib can inhibit the growth of MB tumor *in vivo* and this effect is accompanied by simultaneous reduction in Gli-1 mRNA expression, which is a direct measure of Hh pathway activity.

**Figure 7.**
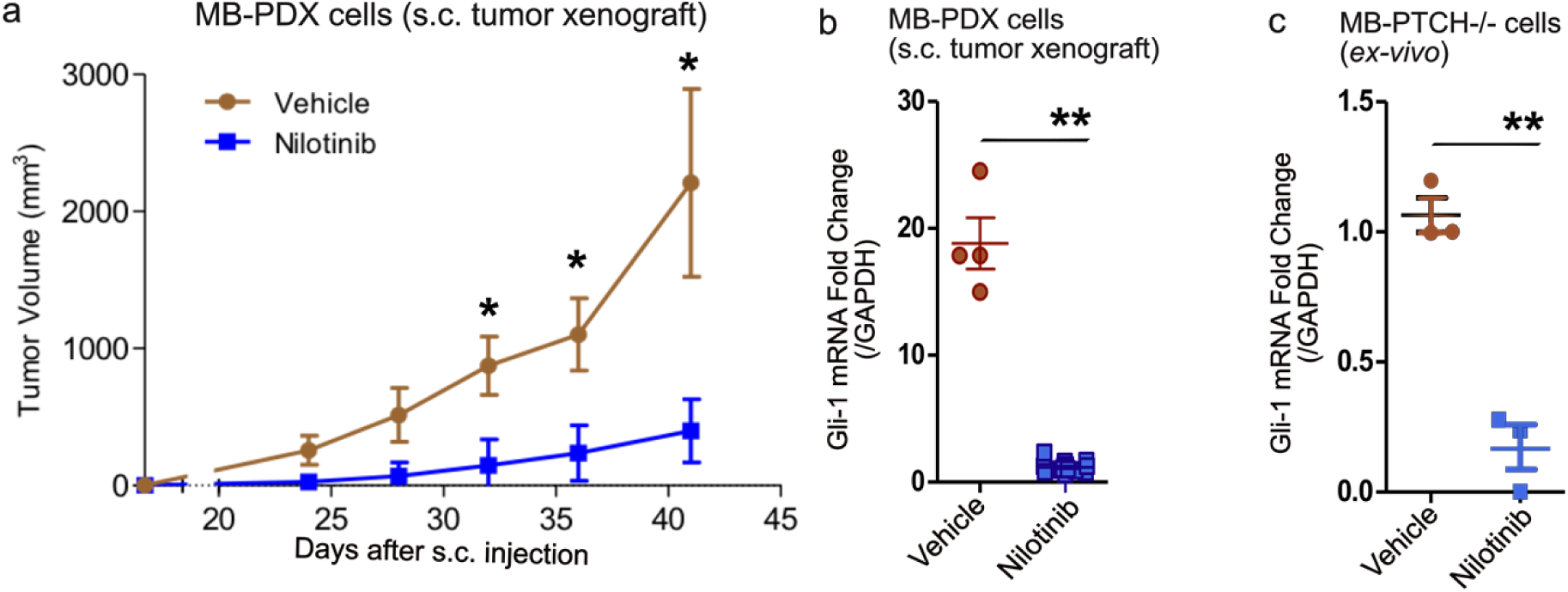
Nilotinib reduces tumor growth and Gli-1 mRNA expression *in-vivo*. **(a)** Nilotinib treatment (40 mg/kg/day) led to significant reduction in tumor volume as compared to vehicle treated group in the MB-PDX xenograft model (n=2). **(b)** Gli-1 mRNA expression is reduced in Nilotinib (10 µM) treated tumors as compared to vehicle treated tumors of MB-PDX xenograft model (n=3). **(c)** Gli-1 mRNA expression is decreased in the MB (Math1-Cre; ptch1-flox/flox) cells propagated in mice and treated *ex-vivo* with Nilotinib (10 µM) in ultra-low attachment plates for 24 hours (n=3). Data represent the mean and standard deviation for repeat experiments, and asterisks represent p-values as follows: *, p < 0.05; **, p < 0.005.

The qRT-PCR data encouraged us to investigate the effect of Nilotinib on Gli-1 mRNA expression in other cell types. We used genetically modified mouse cell in which PTCH1 is removed (Math1-Cre; ptch1-flox/flox cells, referred as PTCH −/−) resulting in constitutively active SMO and Hh pathway and hence pronounced medulloblastoma phenotype^57^. Treatment of these cells with Nilotinib (10 µM) *ex vivo* for 24 hours resulted in significant reduction in Gli-1 mRNA expression as compared to the vehicle treated cells. The results from three independent replicates were used for quantitative analysis and are represented in Figure 7c. These data strongly support the relevance of the Hh pathway inhibition by Nilotinib in medulloblastoma and warrants its further testing in Hh-MB patients.

## Discussion

The critical role of the Hh pathway in several cancers, combined with the limited success of SMO- specific antagonists in MB and other Hh-dependent cancers^19^, motivated us to search for Hh pathway modulators with a different target profile. The desired profile would include SMO *along with* other cancer-related targets because of two considerations: (i) fast emergence of drug resistance with the use of a single specifically targeted drug^58,59^; and (ii) likely increase of toxicity and unfavorable drug-drug interactions with the use of drug combinations^26^. We relied on three assumptions: (i) most approved drugs possess multi-target and multiple pathway modulation at therapeutic concentrations, and these targets are not fully characterized, (ii) some of the yet undiscovered targets of approved drugs may cover complementary relevant pathways, and (iii) the search for the yet unknown targets can be assisted with computational methods.

Discovered primarily as high-affinity specific inhibitors of BCR-ABL fusion protein, Imatinib and Nilotinib have been since shown to have extensive multi-target pharmacological profiles^34^. In addition to their primary target ABL1, both Imatinib and Nilotinib inhibit tyrosine kinases PDGFRα/β and c-Kit^34^. Importantly, both PDGFRα and c-Kit are upregulated in medulloblastoma^60^, and PDGFRβ was shown to be critical for migration and invasion of medulloblastoma cells^61^. Imatinib also inhibits protein kinases DDR1/2, CSF1 and others^62,63^, a total of 12 targets with K_d_ or K_i_ better than 100 nM (a complete list is given in Supplementary Table 2). By comparison, Nilotinib has 18 additional documented targets with K_d_ or K_i_ below 100nM, including protein kinases ABL2, DDR1/2, MK11, MLTK, EPHA8, FRK, LCK, LYN, and several carbonic anhydrases^62,63^ (Supplementary Table 3). Our study demonstrated that in addition to this target list, Nilotinib inhibits Hh signaling with potency and efficacy similar to that of Cyclopamine, and well within the clinically achievable concentration range (Figure 4). Imatinib also inhibited Hh signaling but was substantially weaker than Nilotinib, making its anti-Hh activity clinically irrelevant. Using a competition binding assay, we confirmed the direct binding of Nilotinib and Imatinib to SMO. The results of Western blot analysis and qRT-PCR analysis for Gli-1 protein and mRNA expression, respectively, in MB cells further confirmed that Hh pathway inhibition by Nilotinib is a result of SMO antagonism/inhibition.

Interestingly, this is not the first time when type-II protein kinase inhibitors are found to have specific secondary activity at SMO. In 2012, Yang et al. reported repurposing and optimization of an experimental p38α inhibitor for SMO, although in that case, the optimization led to loss of anti-p38α activity of the compound^64^. Furthermore, a recent report identified Glesatinib and Foretinib (experimental type-II MET tyrosine kinase inhibitors) as negative modulators of SMO using *in-silico* repurposing approach^28^. Taken together, these findings support the concept of the type-II protein kinase inhibitor chemotype being broadly compatible with SMO antagonism and suggest the robustness of concurrent inhibition of several cancer related pathways in SMO-dependent cancers.

The ability to simultaneously inhibit SMO and the specified protein kinases makes Nilotinib a multi-pathway inhibitor with a diverse target profile that includes the Hh pathway (Figure 8). We hypothesized that such multi-pathway inhibition profile may make Nilotinib a superior agent to the SMO-specific antagonists. Corroborating this hypothesis, Nilotinib was observed to be more potent than Vismodegib in reducing cell viability, inhibiting NS formation and nuclear Gli-1 expression in Hh-dependent MB- PDX and DAOY cells. The results of our Western blot and qRT-PCR analysis strongly suggest a direct effect of Nilotinib on the Hh pathway and Gli-1 expression. However, the properties of CSCs self-renewal and survival are controlled by various embryonic signaling pathways including but not limited to the Hh pathway^65^; therefore, the multi-pathway inhibition profile of Nilotinib might have contributed to its cell growth inhibition efficacy. Imatinib was found to be less potent than Nilotinib in these assays, in accordance with the *in vitro* functional and binding experiments.

**Figure 8.**
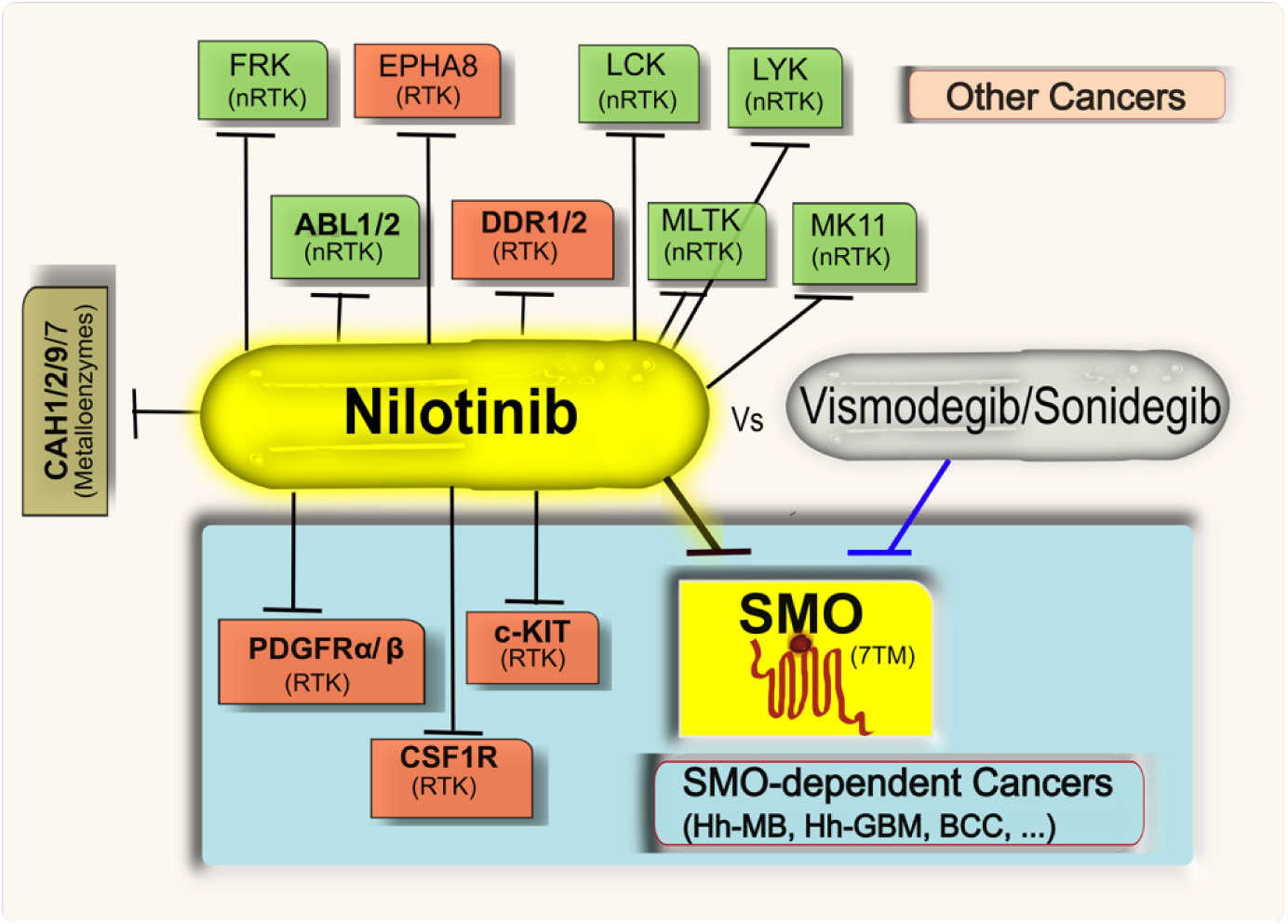
Multi-pathway pharmacology of Nilotinib vs SMO-specific drugs for Hh-dependent cancers. Targets of Nilotinib include the Hh-pathway (identified in this study) and several other pathways that are either already known to be dysregulated in Hh-dependent cancers or may serve as escape pathways and are implicated in other cancers. The ability of Nilotinib to inhibit multiple targets simultaneously in Hh-dependent cancers makes it suitable candidate for therapeutic candidate for personalized medicine as compared to specific SMO inhibitors like Vismodegib. Target type is shown by color. RTK stands for receptor tyrosine kinase and nRTK stands for non-receptor tyrosine kinase.

Because the CSC population is thought to drive cancer relapse and resistance to drugs^12,66^ in patients, the anti-CSC activity of Nilotinib may help overcome these issues. Also, Nilotinib is highly active and safe in Imatinib-resistant patients with chronic myeloid leukemia in chronic phase (CML-CP)^67^. The effectiveness of Nilotinib in CML-CP patients can also be attributed, in part, to its anti-SMO activity reported here and its ability to inhibit CSC growth. In fact, a recent study^68^ demonstrated alterations of Hh pathway gene expression in a subpopulation of CML-CP patients: the negative modulators GLI3 and SUFU genes were down-regulated and the Hh target genes CCNB2, STIL, FOXM1 and GLI1 genes were up-regulated.

Nilotinib is an FDA-approved drug with well-characterized pharmacokinetics and safety profile. It is well tolerated by patients even in the course of long-term treatment^67^. The newly discovered anti-Hh activity of Nilotinib compares favorably with clinically achievable plasma concentrations of the drug in patients, which, given its low toxicity, makes Nilotinib an attractive therapeutic candidate against Hh- dependent cancers, alone or in conjunction with surgery, radiotherapy, and chemotherapy^69,70^. Moreover, despite the initial reports of poor penetration of the blood brain barrier (BBB)^71^, Nilotinib was later shown to be detectable in mouse brains upon oral and intraperitoneal administration^72,73^ and to exhibit pronounced CNS efficacy with long-lasting responses in pre-clinical models of Parkinson’s disease^72^ and in BCR-ABL+ CNS relapse CML patients^71^. These findings suggest that Nilotinib can be used in neurological cancers as well.

To our knowledge Nilotinib has not been specifically studied in the context of Hh-dependent cancers. However, both Nilotinib and Imatinib are being studied for brain cancers that in a fraction of cases are driven by dysregulation of the Hh pathway, e.g. for GBM^10^. In preclinical studies, Nilotinib decreased cell viability and tumorigenicity in GBM cell lines ^47^. Its effects in GBM are currently being investigated in a phase II trial^74^. Imatinib has been tried in GBM patients without much effect on proliferation of cancer but with pronounced effect on biochemical profile of cancer tissue^75^. Finally, pretreatment of a human GBM cell line with Imatinib significantly enhanced cytotoxic effect of ionizing radiation^76^. Taken together with the above considerations, the newly discovered anti-SMO activity of Nilotinib makes it a strong candidate for repurposing to Hh-dependent MB and other Hh-dependent cancers, as well as cancers where the Hh pathway was shown to be activated upon treatment, and warrants its further studies in specific cohorts of Hh-dependent cancer patients.

Discovering new essential targets of existing drugs and matching their real pharmacology with the subtypes of the patients’ tumors, appears especially promising in the context of personalized medicine.

## Materials and methods

### Software

Ligand preparation, preparation of the ligand-based chemical fields, docking, and analyses were carried with ICM versions 3.7-4 and 3.7-6 (Molsoft L.L.C., La Jolla, CA)^77,78^.

### Preparation of SMO complexes

Crystal structures of SMO complexes with LY2940680, SANT-1, Cyclopamine, ANTA XV and SAG (the latter an agonist) were downloaded from the Protein Data Bank (PDB) (http://www.rcsb.org/pdb/home/home.do), with accession codes 4JKV, 4N4W, 4O9R, 4QIM and 4QIN, respectively^29,35^. Tightly bound water molecules were retained and the remaining water molecules were removed. Hydrogen atoms, as well as heavy atoms of missing or disordered residue side chains were rebuilt and optimized. Formal charges were assigned to the ligand molecules at pH 7.4 using the pKa prediction model. The protonation states of D473 and E518 were assigned to neutral to optimize buried polar network in the pocket. After that, the complexes were subjected to restrained conformational minimization (ligands, water molecules, and amino-acid residue side chains only) to eliminate minor clashes and optimize polar interactions between the receptor and the ligands.

### Collection and preparation of chemical dataset

The set of approved and withdrawn drugs from the DrugBank database [http://www.drugbank.ca] was used for screening. Specifically, FDA-approved (1510 as of Nov 2014) and withdrawn (189 as of Nov 2014) small molecule drugs (1699 in total) were downloaded from DrugBank database website. Duplicate compounds, chemical mixtures, polymers, inorganic, and organometallic compounds were removed. Two-dimensional (2D) structures of DrugBank dataset were standardized by removing salts and explicit hydrogens, standardizing chemical group topology, enumerating stereoisomers of racemic compounds, and assigning formal charges at pH 7.4 using the pKa prediction model in ICM. Compounds were converted from 2D to 3D using ICM.

A list of 13 known SMO modulators (one agonist and 12 antagonists, Supplementary Figure 1) was prepared based on literature reports until 2016 and used to estimate the preferred range for logP (1 to 8) and polar surface area (PSA, less than 140). The DrugBank compounds were filtered according to these criteria yielding a set of 720 approved drugs and 128 withdrawn drugs.

### Construction of chemical fields and database screening

The five complexes of SMO with various ligands were superimposed by the vicinity of their binding pockets. The five unique co-crystallized ligands were extracted and used for construction of Ligand-APF (chemical fields) models as described earlier^36^. The co-crystalized ligands occupy two distinct sub-pockets within the binding cavity of SMO, and belong to different chemotypes, and one of the structures (the one with cyclopamine) had a lower resolution of 3.2Å. Hence, separate chemical field models were prepared. The drug database compounds were screened against the chemical field models. For the best compound pose, the chemical field, or APF, score was calculated as a sum of the scores in the seven potential grid maps representing different pharmacophore features as described before^36^.

### Construction of pocket-based models and database screening

The five optimized receptors were converted into potential grid maps representing (i) Van der Waals interaction (calculated as Lennard-Jones potential with hydrogen, carbon, and large-atom probes) (ii) electrostatic potential, (iii) hydrogen bonding potential combining donors and acceptors, and (iv) polar surface energy. The potentials were calculated on a 0.5 Å grid in a box surrounding all crystallographic ligands^79^, with a margin of 2.5 Å on each side. The selected compounds were docked into each of the five sets of maps (representing five different receptor conformations). Prior to sampling, multiple starting poses of each ligand were generated by exhaustively sampling the ligand *in vacuo* and overlaying each of the resulting conformations onto the binding pocket in four principal orientations. The sampling phase was performed using biased probability Monte Carlo optimization of rotational, translational, and torsional variables of the ligand in an attempt to achieve the global minimum of the objective energy function. This energy function is a combination of ligand interactions with the receptor grid potentials and the explicit (full-atom) interactions within the ligand itself (ligand strain).

The five co-crystallized compounds and other SMO ligands (Table 1) were included in the docking procedure as positive controls and to determine the docking score cutoff. Based on their analysis, drugs with predicted docking scores of less than −20 were considered for further experimental testing. Predicted binding poses of selected compounds were visually inspected and literature was consulted for properties and mechanism of action. Fifty (50) top-scoring compounds were inspected visually and checked against literature (Supplementary Table 1). Five drugs were selected for experimental testing in laboratory. Other molecules with favorable docking scores are being further evaluated and will be presented in other publications.

### Chemicals and reagents

Cyclopamine was purchased from Cayman Chemical Company. Imatinib and Nimesulide were purchased from Sigma Life Science. Nilotinib, SAG, Vismodegib and Erlotinib were purchased from SelleckChem. Mebendazole was purchased from Tokyo Chemical Industry (TCI). Thalidomide was purchased from MP Biomedicals. BODIPY-Cyclopamine was purchased from Biovision.

For the assays, test compounds were prepared at 10-fold the final well concentrations in assay media, across six concentrations, using 1:10 or 1:5 serial dilutions from the highest concentration (usually 10 mM). All stock solutions were stored at −20°C.

B27 supplement was purchased from Invitrogen. Human recombinant bFGF and EGF were purchased from Invitrogen/Gibco. Alamar Blue was purchased from Invitrogen. Antibacterial-antimycotic Solution (AAS) was purchased from Gibco.

### Cells and plasmids

HEK293T cells were obtained from ATCC. NIH 3T3 Gli-RE cells (NIH 3T3 stably transfected with firefly luciferase under transcriptional control of 8× Gli response element) were obtained from BPS Bioscience, San Diego, CA. Medulloblastoma Patient Derived Xenograft (MB-PDX) cells were generated by the Durden Laboratory as previously described^51^. Genetic profiling of MB-PDX cells indicates that they originated from the Hh-dependent MB subtype^51^. Math1-Cre; ptch1-flox/flox cells (PTCH -/- cells) were generated by Wechsler-Reya’s lab^57^ and were propagated in NSG mice by orthotopic tumor model. DAOY cells were purchased from ATCC (ATCC^®^HTB-186^™^).

The mSMO and ShhN (Shh N-terminal domain) plasmids purchased from Addgene (#37673 and #37680). All vectors were propagated in XL10 Gold competent cells, purified with NucleoBond Xtra Midi kit (Clontech), and sequenced (Genewiz).

HEK293T cells were cultured in DMEM supplemented with 10% of FBS at 37°C in an atmosphere with 5% CO_2_. NIH 3T3 Gli-RE cells were cultured DMEM supplemented with 10% of BCS and Geneticin. MB-PDX and DAOY cells were cultured in NS culture media [50% DMEM + 50% F12 + 1×B27 supplement + 10 ng/ml bFGF + 10 ng/ml EGF + 1× AAS] or adherent culture media (DMEM + 10% FBS + 1× AAS) in ultralow bind or tissue culture treated flasks, respectively at 37°C in an atmosphere with 5% CO_2_. PTCH1 -/- cells were propagated in *in vivo* conditions (in mouse brain after injecting 1×10^6^ cells per injection/4 µL).

For production of ShhN conditioned media and for the BODIPY-Cyclopamine competition binding assay, HEK293T cells were seeded at the density of 1.5×10^6^ in a 6 cm dish, allowed to grow overnight and then transfected with either ShhN or mSMO plasmid DNA (6 µg DNA per 6 cm dish) using TransIT transfection reagent (Mirus Bio LLC) according to the manufacturer’s instructions. Prior to transfection, cell culture media was replaced with DMEM + 10%FBS for mSMO transfection and with DMEM + 10%BCS for ShhN transfection. The transfected dish was incubated for approximately 24 hours at 37°C. For the production of ShhN conditioned medium, the culture medium from ShhN-transfected HEK293T cells was collected, aliquoted in single use 1.5 mL Eppendorf tubes and either used fresh or stored at −20°C. For BODIPY-Cyclopamine competition binding assay, HEK293T cells transfected with mSMO plasmid were lifted with trypsin (0.25%), re-plated in 96 well adherent plates and incubated for additional 24 hours before the experiment.

### Gli-Luciferase functional assay

NIH 3T3 Gli-RE cell were plated at 1.2×10^4^ cells per well in 100 µL of DMEM+10%BCS media in 96 well tissue culture treated plate (Falcon, 353219). After 24 hours, when the cells reached confluency, the culture medium was replaced with 80 µL per well of assay medium [Optimem + 10 mM HEPES + 1 mM Sodium Pyruvate + 1× MEM NEAA]. Serial dilutions of test compounds (10x of final concentrations) were prepared in assay media. 10 µL of diluted compounds (10x the final concentration) or control media were added to the plate after which the plate was incubated for 15 to 30 minutes at 37°C. Then the cells were stimulated by addition of either ShhN-conditioned media or control media (DMEM+10%BCS) at a final concentration of 10%. Following incubation of the plate for at least 28 hours at 37°C, the cells were simultaneously lysed and supplemented with a luciferase substrate by addition of equal volume of Steady-Glo reagent (Promega, E2520) directly to the assay media. Plates were mixed by pipetting, centrifuged to eliminate foam, incubated for 10 minutes, and analyzed using Perkin Elmer Viktor X luminescence plate reader. The results were analyzed using a non-linear regression (Prism 6, GraphPad Software, La Jolla, CA). Data were normalized to the maximal response observed for ShhN-stimulated cells in the same experiment. A sigmoidal-dose response curve was used as a model for data analysis and IC_50_ value calculation.

### BODIPY-Cyclopamine competition binding assay

HEK293T cells transiently transfected with mSMO were lifted with trypsin (0.25%) and replated at 6×10^4^ cells in 80 µL DMEM + 10%FBS per well in a 96 well tissue-culture treated plate (Falcon, 353219). The plate was incubated at 37°C for 24 hours. Serial dilutions of test compounds (10x final concentrations) were prepared in culture media. 10 µL of diluted compounds or control media were added to the plate after which the plate was incubated at 37°C for 10 min. Next, BODIPY-Cyclopamine was added to each well at the final concentration of 5 nM, except for control wells that were left unstained. Following at least 1.5 hours incubation at 37°C in 5% CO_2_, cells were lifted by vigorous pipetting, transferred to conical bottom 96-well plates, and centrifuged at 400 × g for 5 min at 4°C. Supernatant was discarded, cells were re-suspended in 300 µL of PBS + 0.5% BSA (freshly prepared), and the plate was analyzed with Guava benchtop flow cytometer. The results were interpreted with FlowJo software (version v10.1). Dose response curves were constructed in Prism 6 (GraphPad Software, La Jolla, CA).

### Cell viability assay

MB-PDX or DAOY cells were cultured in NS culture media [50% DMEM + 50% F12 + 1× B27 supplement + 10 ng/ml human recombinant bFGF + 10 ng/ml EGF + 1× AAS] or adherent culture media (DMEM+10% FBS + 1× AAS) in ultra-low attachment (Corning, 3814) or tissue culture treated (Corning, 430641U) flasks, respectively. The cells were plated at 1000 cells per well in 90 µL of respective media in ultralow attachment (Corning, 3474) or tissue culture treated (Falcon, 353219) 96- well plates, and incubated for at least 24 hours. Cells were exposed to test compounds at different serially diluted concentrations (10 µL of 10×final concentration) and again incubated for at least 72 hours. After the incubation period, 5 µL of Alamar blue dye (Invitrogen, DAL1100) was added to each well, the cells were incubated for 2-4 hours, and then analyzed using SpectraMax fluorescence reader using excitation and emission wavelengths of 544 nm and 590 nm, respectively. The results were analyzed using a non-linear regression (Prism 6, GraphPad Software, La Jolla, CA).

### Neurosphere formation assay

MB-PDX or DAOY cells were cultured in NS culture media in ultralow attachment flasks (Corning, 3814). The cells were plated at the density of 1 to 2 cells per well in 100 µL of NS culture media with DMSO or test compounds in ultralow attachment 96 well plate (Corning, 3474). The final concentration of test compounds in the wells was 10 µM. Cells were incubated at 37 °C for at least seven days. After incubation percent of wells positive with neural stem cell-derived sphere colonies were calculated for each test condition and compared.

### Western blot analysis

Nuclear fractions were prepared from cell pellets according to the method of Schreiber *et. al.*^80^ with minor modifications. The MB-PDX or DAOY cells were plated in ultra-low attachment 6-well plates (Costar, 3471) (1×10^6^ cells per treatment condition) in NS formation culture media in the presence of either vehicle or test drugs (10 µM). After 24 hours, cells were lifted from the wells by pipetting, transferred to 1.5 mL Eppendorf tubes, centrifuged at 10,000 × g for 1 minutes at 4°C, and washed once in PBS. Following supernatant removal by vacuum, cells were lysed using a plasma membrane lysis buffer [10 mM HEPES; pH 7.5, 10 mM KCl, 0.1 mM EDTA, 1 mM dithiothreitol (DTT), 0.5% Nonidet P-40 and 0.5 mM PMSF along with 1× protease inhibitor cocktail (Sigma)] and allowed to swell on ice for 15-20 min with intermittent mixing. Next, tubes were vortexed to disrupt cell membranes and then centrifuged at 12,000 × g at 4°C for 10 min. This treatment resulted in separation of cell nuclei as the insoluble fraction. The pelleted nuclei were washed with the plasma membrane lysis buffer and re-suspended in the nuclear extraction buffer [20 mM HEPES (pH 7.5), 400 mM NaCl, 1 mM EDTA, 1 mM DTT, 1 mM PMSF with 1× protease inhibitor cocktail] and incubated on ice for 30 min. The tubes were centrifuged at 12,000 ×g for 25 min at 4°C and the supernatant (now containing the soluble fraction of the cell nuclei) was collected. Total protein concentrations were determined using the Bio-Rad Protein Assay (Bio-Rad, Hercules, CA). Nuclear extracts containing the total of 40 µg protein were separated on a 10% Tris-SDS PAGE gel (Bio-Rad Mini-PROTEAN® TGX). Gels were transferred on nitrocellulose membranes using Bio-Rad TransBlot system, after which the membranes were blocked in the blocking buffer [5% nonfat dry milk in 1× Tris Buffer Saline + 0.1% Tween 20 (TBST)] for 2 hours at room temperature. Following blocking, the membranes were washed four times with TBST (5 minutes gentle rocking per wash) and incubated overnight with primary antibody Rabbit anti-Gli-1 (Cell signaling, #C68H3); 1:250 dilution in TBST-Milk at 4°C with gentle rocking. Membranes were washed three times with TBST for 5 min each. The membranes are incubated with Mouse anti-TBP antibody (BioLegand, #668306), 1:5000 dilution in TBST-milk at room temp. for two hours and then washed three times with TBST for 5 min each. Then membranes were incubated with secondary antibodies (LI-COR Goat anti-Rabbit IRDye-680 and LI_COR Donkey anti-Mouse IRDye-800; 1:10,000 dilution in TBST + 1% BSA) for 1 hour at room temperature. Membranes were again washed three times in TBST and finally transferred into milli-Q water. Membranes were imaged using the Odyssey IR imaging system (LI-COR Bioscience). The intensity of Gli-1 bands was estimated by Image Studio^TM^ Lite (LI-COR Biosciences).

### Subcutaneous Patient-derived Medulloblastoma Xenograft model

All animal studies were performed in accordance with the Animal Care and Use Rules at the University of California San Diego (UCSD). The protocol was approved by Institutional Animal Care and Use Committee (IACUC) of UCSD. NSG mice, 4-6 weeks old, were injected subcutaneously at shoulders and flanks with 1 × 10^6^ viable MB-PDX cells in 1:1 HBSS and gelatinous protein mixture Matrigel. After seven days, vehicle or Nilotinib (40 mg per Kg) treatment was started and given daily intraperitoneally. The tumor size was measured three times a week with a caliper, and tumor volumes were calculated according to the formula (length × width × width × 0.5). At day 42 post-grafting, all mice were euthanized by isoflurane inhalation and cervical dislocation. Tumors were harvested and stored for qRT-PCR analysis.

### Intracranial injection and tumor preparation

An *in vivo* medulloblastoma model was established as described before^57^. Briefly, 6-7 weeks old NOD-scid IL-2Rg null (NSG) mice (the Jackson Laboratory) were anesthetized with xylazine-ketamine, and right carotid artery was exposed. Dissociated Math1-Cre; ptch1-flox/flox cells (1×10^6^ cells in 4 µL HBSS)^57^ were injected into the mouse brains using murine stereotaxic system (Stoelting Co). The coordinates were: 1.8mm to the right of bregma and 3mm deep from the dura. To disassociate the orthotopic tumor into the single cells for the serial implantation or the *ex-vivo* experiment, the tumor was dissected from the cerebellum, followed by digestion using Papain (10 U per mL), DNase (250 U) and L-cystein (2 mg) at 37°C for 30 min.

### Real-time PCR

Both post-vivo MB-PDX cells and Math1-Cre; ptch1-flox/flox cells were analyzed for Gli-1 mRNA expression. For that, total RNA was extracted from dissociated single cells using QIAGEN RNease Mini Kit and cDNA was prepared using 1µg RNA with iScript cDNA Synthesis Kit (Bio-Rad). SYBR green-based QPCR was performed using murine primers listed as below (GLI1, Forward 5’- CCAAGCCAACTTTATGTCAGGG-3’, Reverse 5’-AGCCCGCTTCTTTGTTAATTTGA-3’, GAPDH Forward 5’-ACCCAGAAGACTGTGGATGG-3’, Reverse 5’-TTCTAGACGGCAGGTCAGGT-3’). mRNA levels were normalized to GAPDH (dCt = Ct gene of interest – Ct GAPDH) and reported as relative mRNA expression (ddCt = 2-^(dCt sample – dCt control)^) in fold change.

### Statistical Analysis

Data were expressed as mean ± standard deviation for all experiments. Quantitative analysis of band intensities was performed with ImageStudio, and the data were transferred to GraphPad Prism 7.0 for analysis. Statistical analyses were performed using Welch’s test (an adaptation of student’s t-test) for comparing two groups using GraphPad Prism 7.0.

## Acknowledgements

Authors would like to thank Alok Singh and Muamera Zulcic (Durden Lab, UCSD Moores Cancer Center) for useful discussions and for providing insights for experiments involving MB-PDX cell line, Handel’s lab (UCSD Skaggs School of Pharmacy) for generously providing access to lab equipment, Andrey Ilatovskiy (UCSD Skaggs School of Pharmacy) for helping with the Pocketome tools and pocket comparison, Siamak Amirfakhri (UCSD Moores Cancer Center) for helping in handing of animals, Arash Garossian (UCSD Skaggs School of Pharmacy) for stimulating discussions at the early stages of the project, John R. Crawford (UCSD Department of Neurosciences) for useful discussions.

## Supplementary data (provided as separate files)

**Supplementary Figure 1:** Calculated properties of SMO Modulators and Drugs Selected for Experimental Validation.

**Supplementary Figure 2:** Effect of test drugs on Hh pathway activity with and without ShhN

**Supplementary Figure 3:** Nilotinib, Imatinib, and Vismodegib inhibit luciferase reporter activity in an Hh-dependent manner.

**Supplementary Figure 4:** Original Western Blot images presented as Figure 6(f) in the manuscript for quantitation of Gli-1 (MW – 150) after treatment with DMSO or 5 μM Nilotinib/Vismodegib for 24 hours in MB-PDX and DAOY cells using Odyssey IR imaging system.

**Supplementary Table 1:** List of Top Scoring Molecules in SMO Docking

**Supplementary Table 2:** List of Imatinib Targets and the Kd/Ki values

**Supplementary Table 3:** List of Nilotinib Targets and the Kd/Ki values

